# Dorsolateral prefrontal cortex TMS evokes responses in the subgenual anterior cingulate cortex: Evidence from human intracranial EEG

**DOI:** 10.1101/2024.12.20.629857

**Authors:** Nicholas T. Trapp, Xianqing Liu, Zhuoran Li, Joel Bruss, Corey J. Keller, Aaron D. Boes, Jing Jiang

## Abstract

Transcranial magnetic stimulation combined with intracranial local field potential recordings in humans (TMS-iEEG) represents a new method for investigating electrophysiologic effects of TMS with spatiotemporal precision. We applied TMS-iEEG to the dorsolateral prefrontal cortex (dlPFC) in two subjects and demonstrate evoked activity in the subgenual anterior cingulate cortex (sgACC). This study provides direct electrophysiologic evidence that dlPFC TMS, as targeted for depression treatment, can modulate brain activity in the sgACC.

## 1. Introduction

The subgenual anterior cingulate cortex (sgACC) is implicated in the pathophysiology of various psychiatric disorders, particularly major depressive disorder (MDD)^1–3^. PET imaging studies suggest that sgACC activity correlates with depression severity, and effective antidepressant treatment tends to reduce sgACC activity across a range of interventions^2^. As a result, this region has been a target for neurostimulation procedures in psychiatry, including deep brain stimulation^1,4^ and transcranial magnetic stimulation (TMS)^5^.

TMS is a noninvasive neurostimulation tool that induces neuronal activity changes via electromagnetic induction^6^. Repetitive TMS delivered to the dorsolateral prefrontal cortex (dlPFC), another brain region with functional importance in the pathophysiology of depression^7^, has demonstrated efficacy in treating major depressive disorder (MDD) across several randomized, sham-controlled trials and meta-analyses^8^. One hypothesis is that therapeutic stimulation of the dlPFC with TMS has antidepressant effects through engagement of the sgACC. The sgACC is too deep for direct modulation using current TMS coils, but sgACC modulation via dlPFC stimulation could be possible if the effects of TMS propagate to the sgACC. Whether this remote sgACC modulation occurs is an open question. The left dlPFC is often functionally anticorrelated (inversely correlated) with the sgACC, and evidence suggests the strength of this anticorrelation is associated with treatment response^9^. Indeed, this dlPFC targeting strategy represents the first FDA-cleared functional connectivity MRI (fcMRI)-guided TMS treatment protocol^5,9^.

It is unclear whether dlPFC TMS can modulate sgACC activity, yet this topic has imminent clinical implications in light of these currently-used fcMRI clinical targeting strategies^10–12^. Primate tracer studies suggest sparse or a complete lack of direct structural connections between non-human primate equivalents of the dlPFC and sgACC^13,14^. Recent studies of this relationship with concurrent TMS-fMRI demonstrate TMS-evoked BOLD responses in the sgACC following DLPFC stimulation^15–17^. However, assessing sgACC responses with fMRI remains contentious due to the sluggishness of the BOLD response and the sgACC region’s physical proximity to artifact-generating anatomy (e.g., sinus cavities) that can result in low signal-to-noise ratios. Scalp EEG signals offer better temporal resolution but provide limited knowledge of focal sgACC activity due to the inherent limitations of source localization^18^.

To address this knowledge gap, we utilized a new method that combines TMS with intracranial EEG recordings (TMS-iEEG), which has greater spatial and temporal resolution to identify the presence of evoked responses in specific anatomical locations^19–21^. We show data from two subjects who had single-pulse TMS applied to the dlPFC, targeted to the same region as clinically indicated for the treatment of depression^22,23^, while simultaneously recording from iEEG depth electrodes implanted within the sgACC.

## 2. Results

Among the twelve enrolled subjects who received single-pulse neuronavigation-guided TMS to the dlPFC, we identified two subjects with iEEG depth electrode coverage within the sgACC. One subject (Patient 460) received TMS to left dlPFC (**Fig. 1a)**; the other subject (Patient 625) received TMS to the right dlPFC (**Fig. 1c)** due to anchor bolts associated with the implanted electrodes impeding access of the TMS coil to the left frontal lobe (**Supplementary Fig. S1f**). Each subject had one electrode contact within the left sgACC (**Fig. 1b,d,e,f**). We observed significant intracranial TMS-Evoked Potentials (iTEPs) recorded from the sgACC contacts in both subjects (**Fig. 1g,h; Supplementary Tables S1, S2**). Specifically, amplitudes of iTEPs at both sgACC contacts following active TMS exceeded 5 standard deviations (SDs) within the time window [25 ms, 500 ms] compared to the baseline period [-200 ms, -25 ms], whereas iTEPs following sham TMS did not. Furthermore, the iTEP amplitudes of both contacts following active TMS were significantly higher than those following sham TMS through a nonparametric clustering analysis of the time window 25 to 500 ms post-stimulus (*p* < 0.05, 1000 permutations; for a full overview of the iTEP significance of all contacts in these two subjects, refer to **Supplementary Fig. S1e,f** and **Supplementary Tables S1-2**).

**Figure 1.**
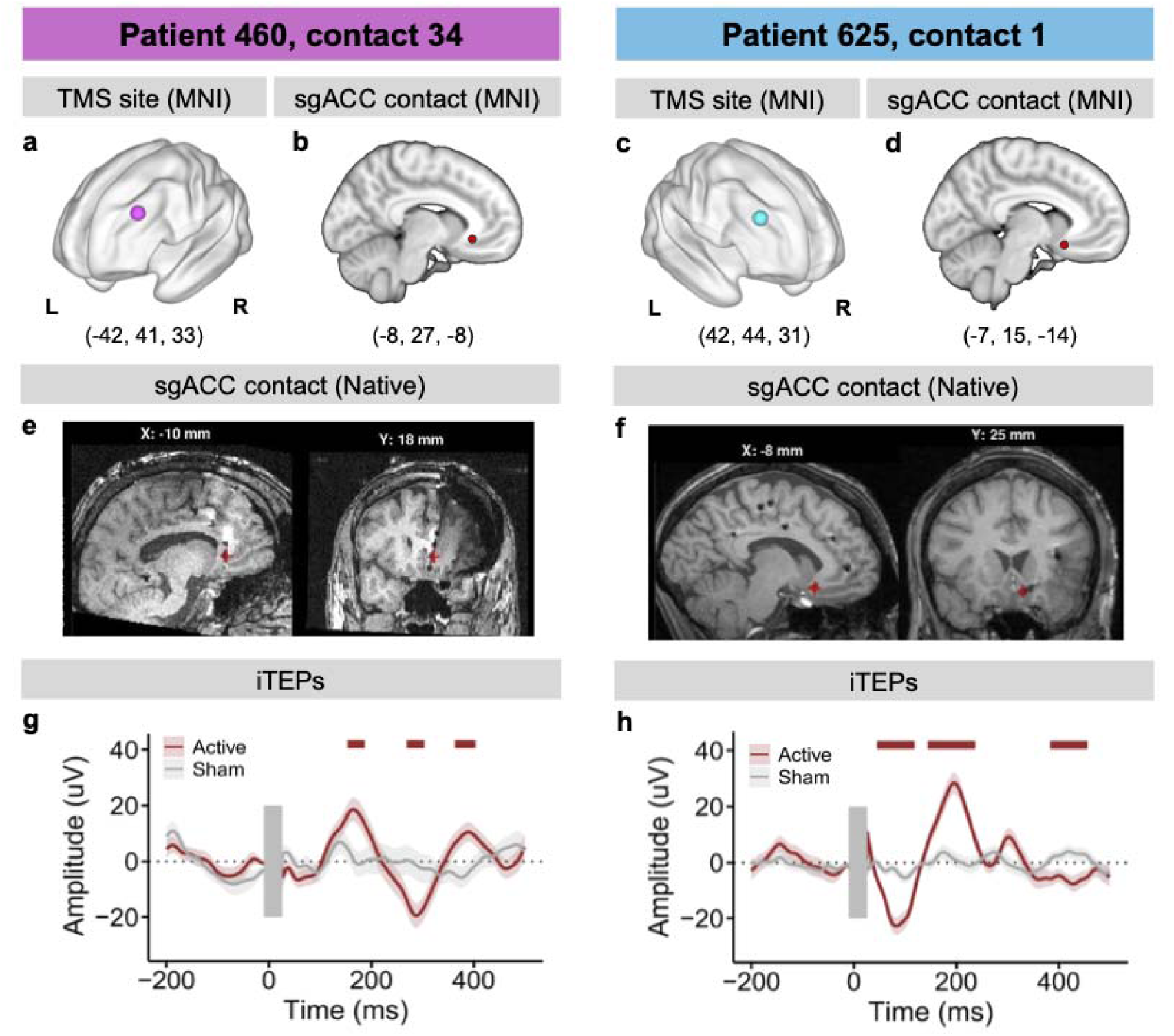
iTEPs in the sgACC following TMS to the dlPFC. (**a, c)** dlPFC TMS sites in MNI space, (**b, d)** sgACC contacts in MNI space, (**e, f)** sgACC contacts in native space**, (g, h)** iTEPs recorded with sgACC contacts in Patient 460 (left panel) and Patient 625 (right panel), horizontal bars marked the first three clusters showing significant active vs. sham iTEP differences (*p* < 0.05) through a nonparametric clustering method with 1000 permutations. dlPFC, dorsolateral prefrontal cortex; sgACC, subgenual anterior cingulate cortex; iTEPs, intracranial TMS-evoked potentials.

In evaluating the specificity of the iTEP, we did not observe significant iTEPs at neighboring regions with coverage in patient 460, such as electrodes on the same depth electrode shaft (1 cm away) and on a separate nearby depth electrode (2.2 cm away, **Supplementary Fig. S2 and Supplementary Table S1**). Subject 625 did not have any nearby electrode coverage in frontal gray matter, although there were no significant iTEPs in three nearby amygdala electrodes (2.6 to 3.1 cm away from sgACC, see **Supplementary Table S2**, electrodes 130-132, *p* = 0.13 to 0.41). To demonstrate the observed effect was not caused by TMS-accompanied auditory or somatosensory artifacts, we also delivered single pulses of intracranial electrical stimulation (iES) through two electrocorticography contacts at a nearby dlPFC region (1.4 cm from the TMS site, 4.5 mA x 60 biphasic pulses jittered at 0.5 Hz, 200 µs pulse width) and observed a significant cortico-cortical evoked response in the sgACC electrode (**Supplementary Fig. S3**) despite having no auditory or somatosensory component to the iES.

We further examined the resting-state functional connectivity (FC) between the dlPFC TMS site and the sgACC contact using both group-level data from a healthy cohort of adults (N=1000) and using individualized, single-subject data, as both have been used to guide treatment targeting^5,9^. The group-population FC (i.e., normative FC) was derived using the Genomics Superstruct Project, GSP, with 1000 healthy participants (18–35 years old, mean age 21.3 years, 50% men)^24^. The single-subject FC was based on subject’s own resting-state fMRI. The normative FC between the stimulation site and the sgACC electrodes showed anticorrelation (Patient 460: *r* = -0.24, *T* = -35.72; Patient 625: *r* = -0.15, *T* = -25.30) across the GSP 1000 participants (**Fig. 2a-b**). Interestingly, the FC between the dlPFC TMS site and the sgACC contact was weaker and variable between the two patients using individual subject-level data (Patient 460: *r* = -0.16; Patient 625: *r* = -0.01) (**Fig. 2c-d**). Whole brain functional connectivity with the dlPFC stimulation sites as the seed regions of interest, as opposed to the sgACC as the seed, are included in **Supplementary Fig. S1a-d** for completeness.

**Figure 2.**
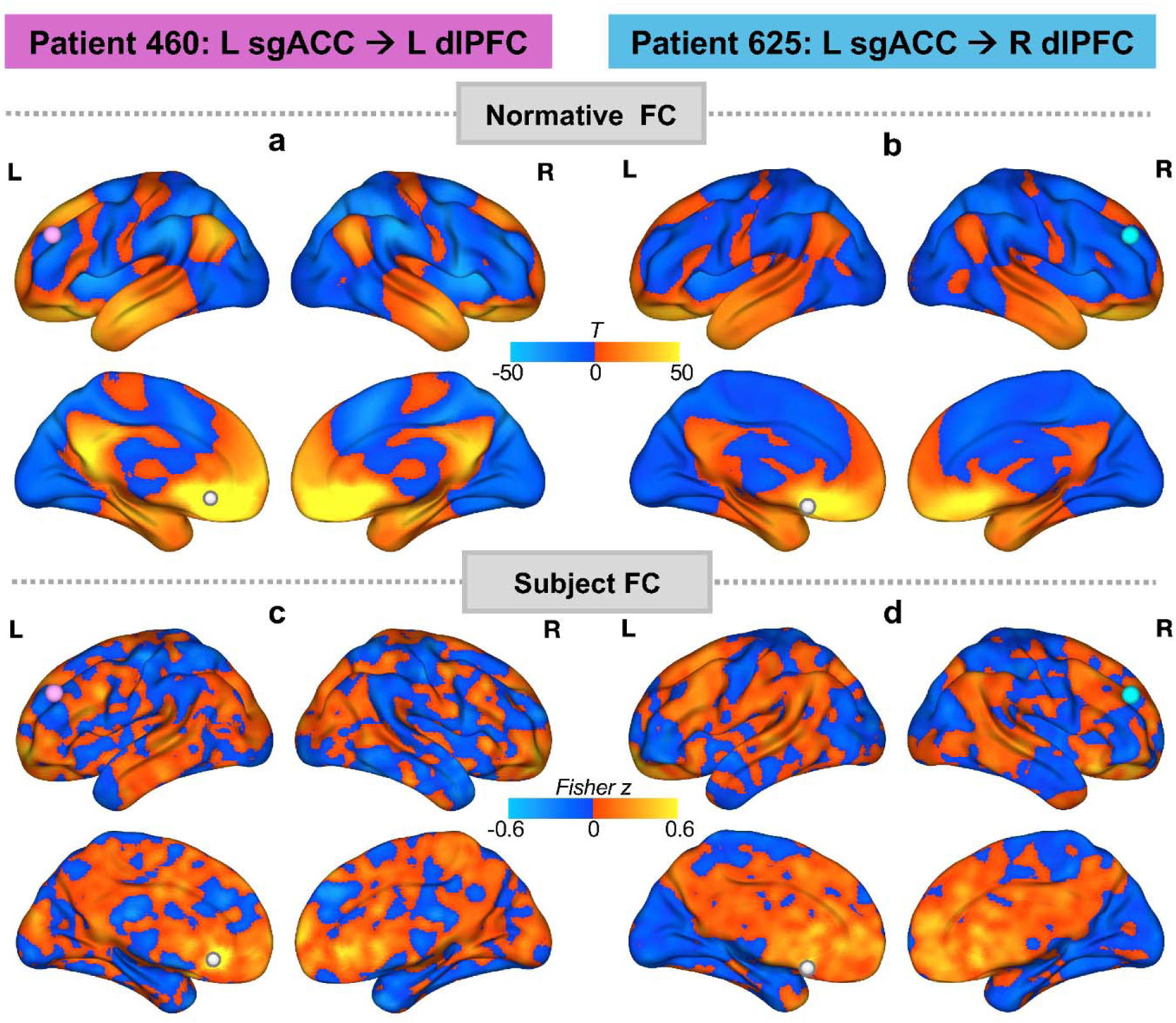
Whole-brain FC maps seeded at sgACC contact. (**a-b)** Normative FC map based on a normative connectome database (*N*=1000); (**c-d**) Subject FC map based on subject’s own resting-state fMRI. sgACC contact as the seed region (white node), TMS sites highlighted with magenta and cyan nodes.

## 3. Discussion

TMS-iEEG represents a new method for investigating the electrophysiologic effects of TMS with greater spatiotemporal precision than previously available with TMS and scalp EEG or fMRI. Here we demonstrate in two subjects, as a proof of principle, that TMS applied to the dlPFC can evoke activity in the sgACC. In doing so, this study provides some of the first direct electrophysiologic evidence in the human brain that TMS of the dlPFC, as targeted for antidepressant treatment protocols, can modulate brain activity in the sgACC. This data further supports the notion that dlPFC TMS generates polysynaptic electrophysiologic responses which can be remote from the stimulation site^19,20^; the evoked responses in the sgACC with direct electrical stimulation of the dlPFC in subject 460 suggest the sgACC response is not due to pain, auditory, or somatosensory experiences. The results also demonstrate that the effects of TMS can occur in brain regions and networks with inverse functional connectivity to the stimulated cortical region. Taken together, this information lends credence to the hypothesis postulating that dlPFC-to-sgACC effects could have a role in mediating antidepressant treatment responses observed with TMS.

There are several limitations to this work. First, TMS-related electrical artifact makes it impractical to look at evoked activity earlier than ∼25 ms post-stimulus with the current approach, which precludes evaluation of the very early responses that may be seen with monosynaptic or oligosynaptic responses^25^. Second, no other “control” sites were stimulated in this experiment to demonstrate the site-specificity of the sgACC iTEPs. Third, this study focuses on single pulse TMS protocols and is thus unable to further describe the modulatory effects of typical repetitive TMS protocols. Finally, these studies took place in patients with epilepsy. Although the sgACC electrodes were not seizure foci nor showing epileptiform activity, it is possible that patients without epilepsy may have different brain responses to TMS in these regions; it is also possible that individual patient factors may lead to unique brain responses to TMS in these regions.

## Supplementary Materials

**Figure S1.**
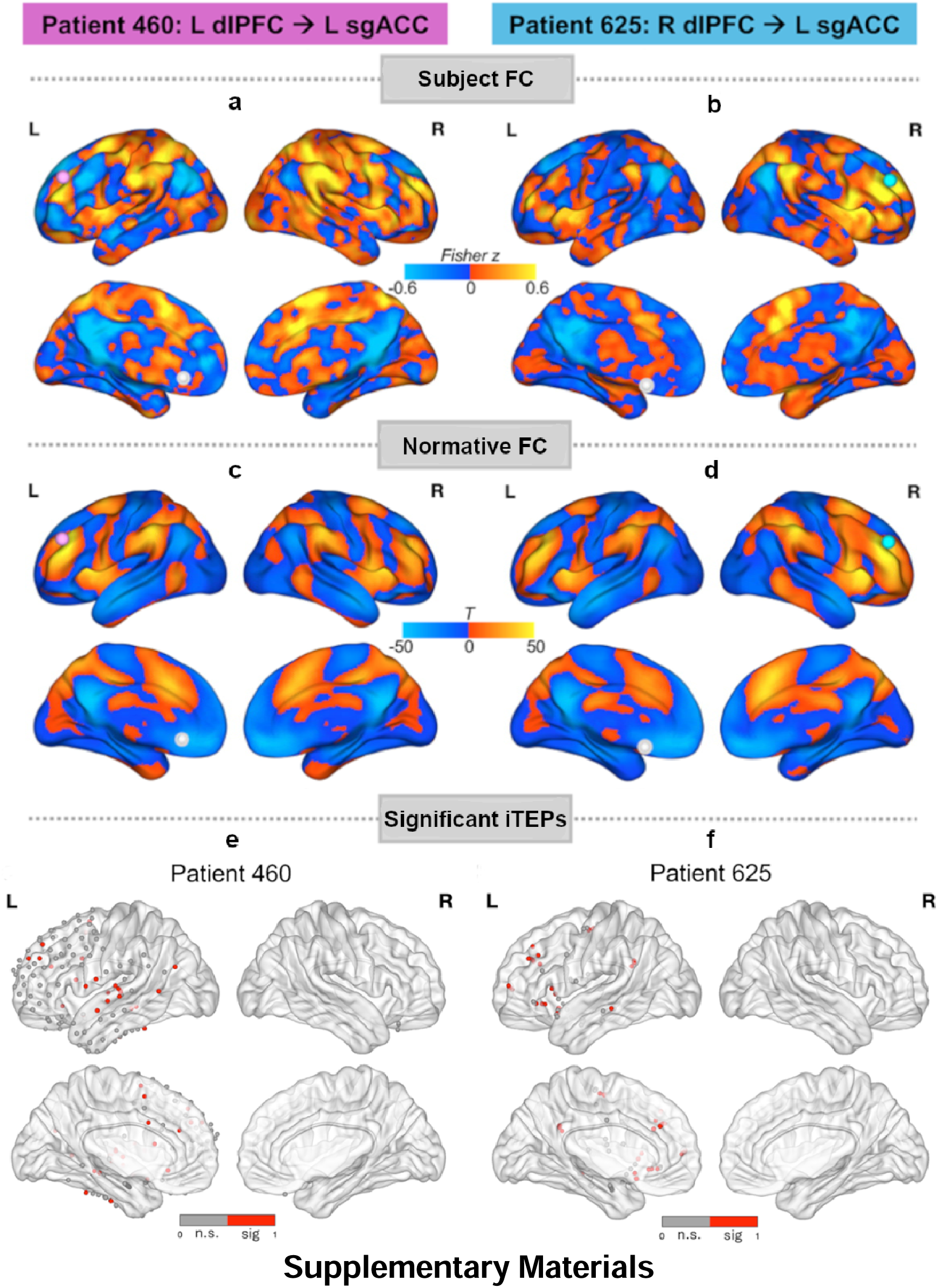
FC between dlPFC TMS site and sgACC contact, and associated significant iTEP findings. (**a-b)** Subject FC map based on subject’s own resting-state fMRI, with TMS site as the seed region (magenta and cyan nodes), and sgACC contact denoted as white nodes. (**c-d**) Normative FC map based on a normative connectome database (*N*=1000), using TMS stimulation site as the seed region of interest. (**e-f**) MNI locations and significance results of whole-brain electrode contacts in Patient 460 (e) and Patient 625 (f). Sig, significant; n.s., not significant. For more details, see **Supplementary Table S1-2**.

**Figure S2.**
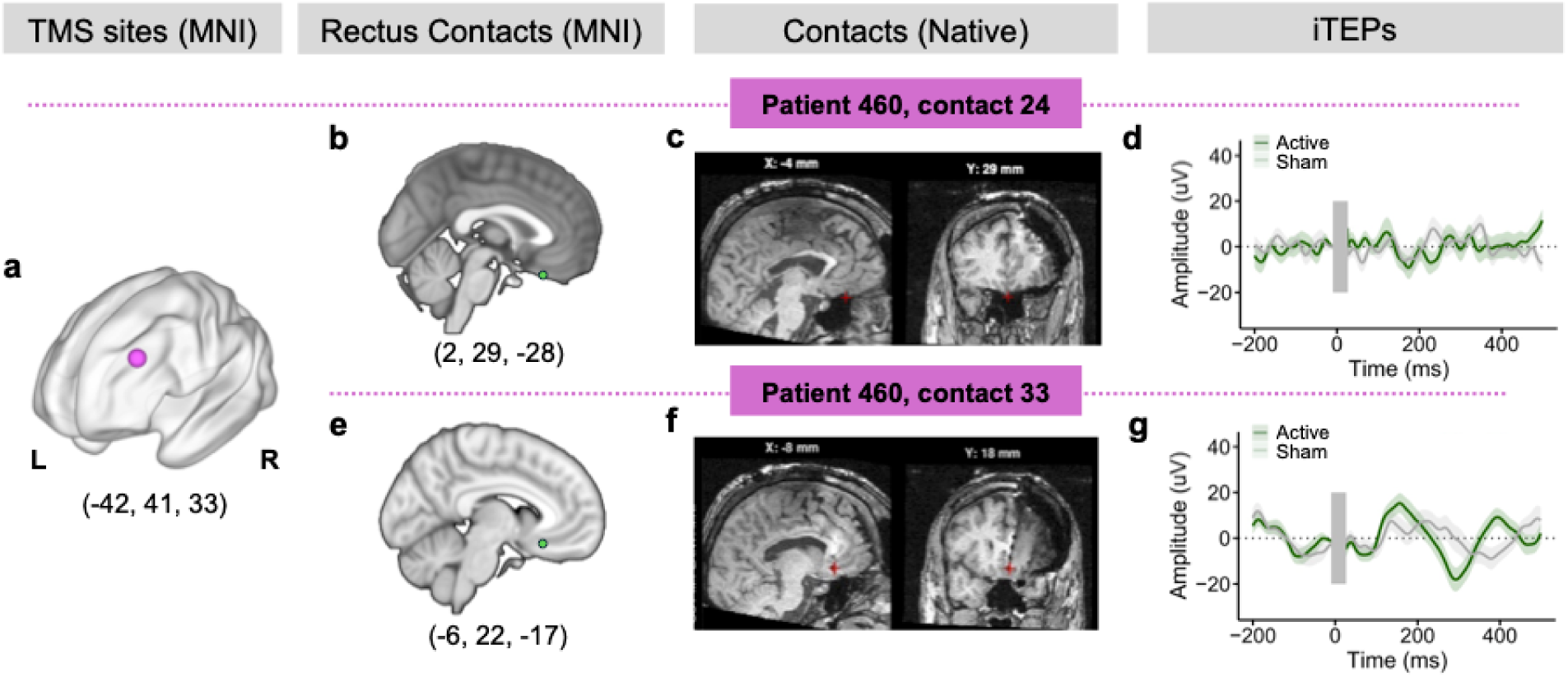
iTEPs in the gyrus rectus following TMS to the dlPFC. **(a)** dlPFC TMS site in MNI space. **(b, e)** Gyrus rectus contacts in MNI space in one of the subjects with sgACC coverage, including one contact on a nearby depth electrode **(b)** and one on the same depth electrode but distal to the sgACC contact **(e). (c, f)** Gyrus rectus contacts in native space**. (d, g)** iTEPs recorded with gyrus rectus contacts in Patient 460; no significant active vs. sham iTEP differences (*p* > 0.05) using a nonparametric clustering method with 1000 permutations. dlPFC, dorsolateral prefrontal cortex; iTEPs, intracranial TMS evoked potentials. Numbers in parentheses refer to MNI coordinates of the region of interest.

**Figure S3.**
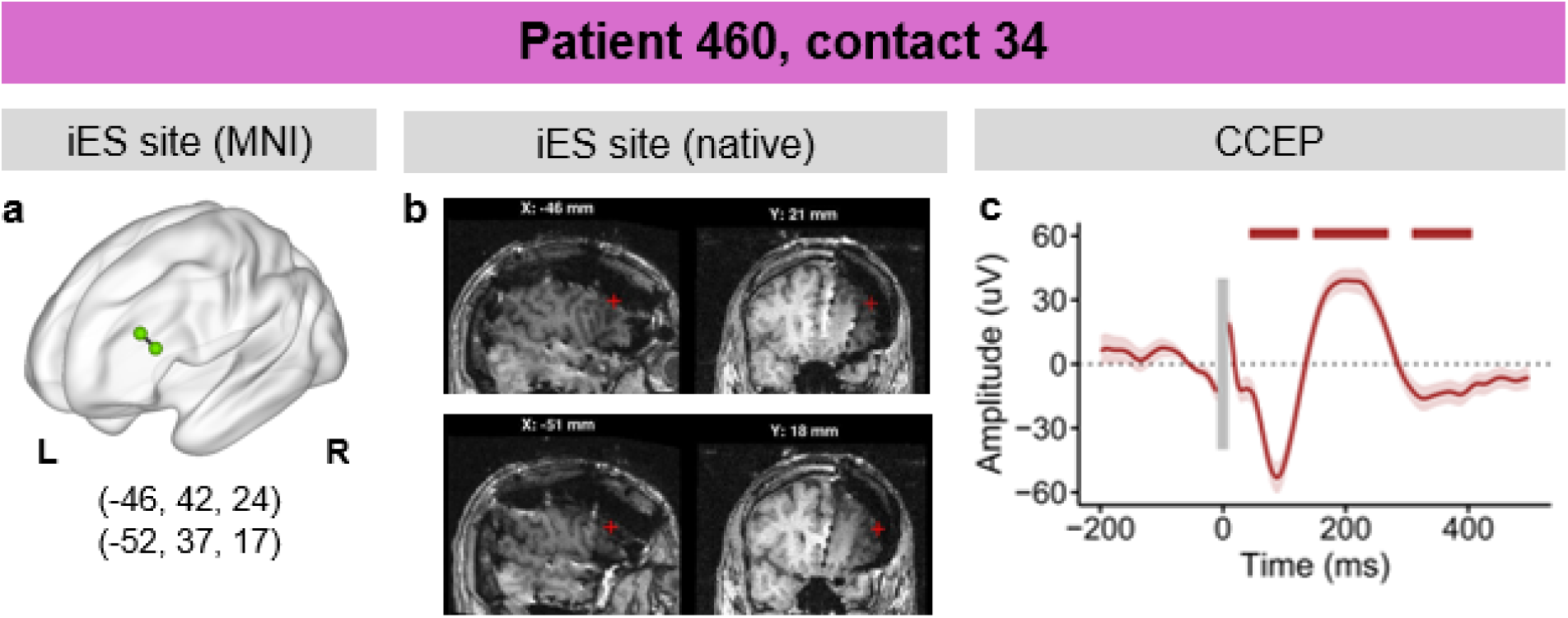
CCEPs in the sgACC following iES to the dlPFC. **(a)** dlPFC iES sites in MNI space, **(b)** iES contacts in native space, **(c)** CCEP recorded with sgACC contacts in Patient 460, horizontal bars mark the three clusters showing significant amplitude change from baseline (*p* < 0.05, FDR corrected). dlPFC, dorsolateral prefrontal cortex; sgACC, subgenual anterior cingulate cortex; iES, intracranial electrical stimulation; CCEP, cortico-cortical evoked potential.

**Figure S4.**
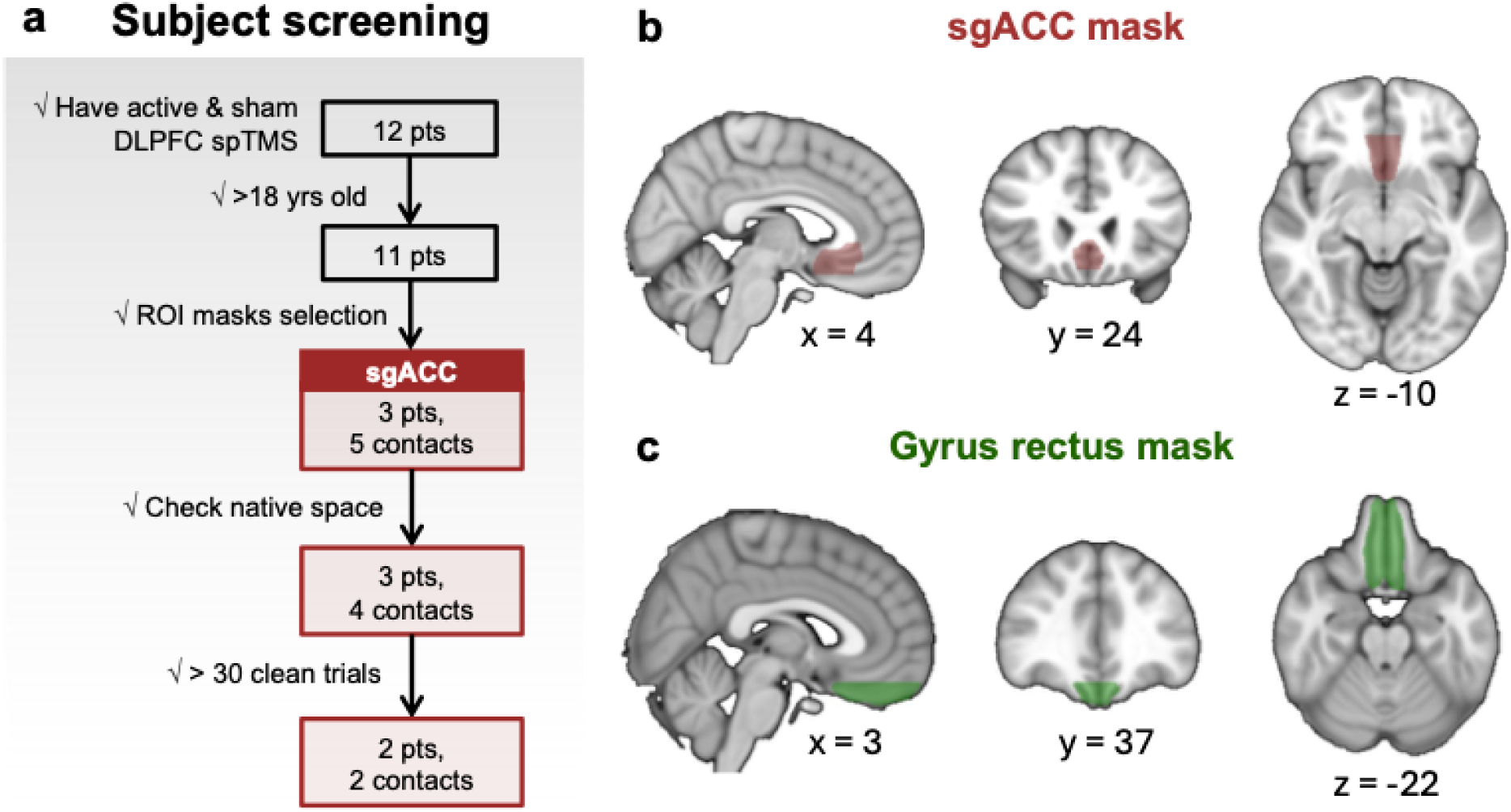
Subject Screening and ROI masks. **(a)** Subject Screening criteria and procedure. **(b)** sgACC mask, which combines the subcallosal cingulate cortex (SCC) mask from the Harvard-Oxford atlas (1mm, 25% threshold) with the sgACC region from the Automated Anatomical Labeling 3 (AAL3) atlas (1mm). Voxels below z = -18.5 (approximate boundary between the SCC and the gyrus rectus) and those within olfactory sulcus were excluded. (**c**) Gyrus rectus mask; voxels above z = -18.5 were excluded.

## Methods

### Participants

For the present study, we screened twelve subjects with medically intractable epilepsy who had consented to participate and were enrolled in a larger TMS-iEEG study as approved by the University of Iowa Institutional Review Board. Criteria for inclusion were age >18 and iEEG electrode coverage in the subgenual anterior cingulate cortex (sgACC) region as part of their clinically indicated pre-surgical epilepsy monitoring. The screening process is shown in **Fig. S4.** Two subjects met inclusion criteria. Subject 1 (460) was a mid-50s male and subject 2 (625) was a mid-20s female. Both were right-handed, had epilepsy onset in their teens, had left mesial temporal lobe seizure foci, and had at least 12 years of formal education. Subject 1 had prior psychiatric diagnoses of major depressive disorder and generalized anxiety disorder; subject 2 had no psychiatric diagnoses as evaluated with a Mini-International Neuropsychiatric Interview^26^.

### Experimental design

Patients underwent structural and resting-state functional MRI (rs-fMRI) prior to their intracranial electrode implantation. Patients underwent a second structural MRI and thin-slice volumetric computerized tomography (CT) for clinical purposes and electrode contact localization on the day following implantation surgery. The TMS-iEEG experiment was conducted after seizures were localized, 1-2 days before electrode explantation and 24 hours after restarting anti-seizure medications. Prior work has demonstrated the safety and feasibility of this experimental method^19^.

### Pre- and post-implantation imaging scans

Pre-implantation structural MRI was acquired using a 3T GE Discovery 750 W scanner with a 32-channel head coil. A 3D FSPGR BRAVO T1 weighted (T1w) sequence was used: TRlJ=lJ8.49 ms, TElJ=lJ3.28 ms, TIlJ=lJ450 ms, FOVlJ=lJ25.6 cm, Flip anglelJ=lJ12°, voxel size = 1.0lJ×lJ1.0lJ×lJ0.8lJmm. The rs-fMRI scan was acquired as five 5-minute gradient-echo EPI runs (650 volumes): TR = 2260 ms, TE = 30 ms, FOV = 22.0 cm, Flip anglelJ=lJ53°, 60 slices (2.5 mm thick, voxel size = 3.44 x 3.44 x 4 mm) covering the whole brain. Post-implantation MRI was also acquired for each patient using a 3T Siemens Skyra scanner with a head transmit-receive coil. An MPRAGE sequence was used: TRlJ=lJ1970 ms, TElJ=lJ3.44 ms, TIlJ=lJ1000 ms, Flip anglelJ=lJ10°, FOVlJ=lJ25lJcm, voxel size = 1.0lJ×lJ1.0lJ×lJ0.8lJmm. The CT acquisition voxel size was 0.47 x 0.47 x 1.0 mm.

### MRI processing

Standard MRI preprocessing steps were performed with the pre-implantation scans using the fMRIPrep pipeline^27^. The T1w image was processed with ANTs^28^ for intensity nonuniformity correction, skull-stripping, and spatial normalization to the MNI template through nonlinear registration. Cerebrospinal fluid (CSF), white-matter (WM) and gray-matter (GM) were segmented on the brain-extracted T1w image using FAST (FSL). For each rs-fMRI run, preprocessing steps included removal of the first 6 volumes, head-motion estimation using MCFLIRT (FSL), co-registration to the T1w reference using boundary-based registration with FLIRT (FSL), distortion correction using a blip-up/blip-down “phase difference” fieldmap acquisition, and resampling onto the standard MNI space. Confounding variables such as framewise displacement (FD)^29^, DVARS and three global signals from the CSF, the WM, and the whole brain were also estimated.

Post-processing steps were performed using XCP-D^30,31^. First, 36 confounding regressors were selected from fMRIPrep outputs. They included 6 motion parameters, 3 mean signals from the CSF, the WM, and the whole brain, temporal derivatives of these 9 parameters, and the quadratic expansion of both the 9 parameters and their temporal derivatives. Next, high-motion outlier volumes were identified with FD value > 0.5 mm and used for data interpolation. The interpolated data and confounds were then detrended and mean-centered, followed by 0.008-0.08 Hz bandpass filtering. The filtered confounds were then regressed from the filtered BOLD data. Finally, the resulting data were censored (high motion volumes removal) and smoothed with a 6-mm FWHM Gaussian kernel.

### TMS site localization

DLPFC stimulation targets were defined using a previously published left dlPFC MNI coordinate (-41.5, 41.1, 33.4) and its contralateral homologue^22,23^ found to be in close proximity to the Beam F3/4 region^32^. The stimulation site was modified slightly if access was impeded by head wrap or anchor bolts. The dlPFC TMS stimulation sites are shown in **Fig. 1a,c**.

### Concurrent TMS and iEEG recording

A Cool-B65 Active/Placebo (A/P) liquid-cooled butterfly coil with a MagVita X100 230V stimulator system (Magventure, Alpharetta, GA, USA) was used for delivering TMS pulses. Brainsight Neuronavigation (Rogue Research, Montreal, Quebec, Canada) with frameless stereotaxy was used to guide stimulation to targets identified on the pre-implantation structural MRI images. Prior to the experiment, motor threshold (MT) was defined for each patient as the stimulation intensity that induced visually observable hand movements in at least 3 of 5 consecutive trials by stimulating the hand knob of primary motor cortex.

During the TMS-iEEG experiment, 50 single biphasic pulses of TMS at 0.5 Hz were delivered to the dlPFC defined above. Stimulation intensity was applied at 100% MT. As a control condition, 50 sham pulses were delivered to the same target with the coil flipped 180°, directed away from the head. Earplugs were used in both conditions to reduce the influence of auditory confounds. During both conditions, iEEG data acquisition was simultaneously made by a multichannel data acquisition system (ATLAS, Neuralynx, Tucson, AZ) with a reference contact placed in the patients’ subdural space. The sampling rate was 8 kHz with 1–2000 Hz acquisition filters (−6lJdB, 256 tap length).

### Intracranial electrode implantation and localization

Patients were implanted with depth electrodes and/or subdural grid/strip arrays (Ad-Tech Medical, Racine, WI). Each depth electrode shaft consisted of 4 to 10 platinum macro-electrode contacts (1.3 mm diameter, 1.6 mm length, 5–10 mm inter-electrode distance). Subdural arrays were flexible silicon sheets embedded with platinum-iridium disc-shaped contacts (2.3 mm diameter, 5–10 mm inter-contact distance).

To precisely determine contact locations for each patient, the post-implantation MRI and CT imaging was linearly co-registered to pre-implantation MR scans using FLIRT (FSL). Images were carefully corrected for displacement and distortion caused by electrode implantation using non-linear 3D thin-plate spline warping with 50-100 manually selected control points. Coordinates of each contact identified on the native space of patients’ post-implantation MRI and CT scans were thus projected onto the pre-implantation MRI space and then normalized to standard MNI (ICBM152 template) space using ANTS^28^.

### sgACC ROI generation

A sgACC mask in MNI space was used to identify the contacts that fall within this region of interest (ROI). This sgACC mask (**Supplementary Fig. S4b**) was created by combining the subcallosal cingulate cortex (SCC) from the Harvard-Oxford atlas (1 mm, 25% threshold)^33^ with the sgACC region from the Automated Anatomical Labeling version 3 (AAL3) atlas (1 mm)^34^, and removing voxels that are below z = -18.5 (approximate boundary between the SCC and the rectus gyrus) or within olfactory sulcus. We visually reviewed the location of each selected contact in its native space to confirm its location within the sgACC.

### iEEG preprocessing

The preprocessing steps were conducted using a customized MATLAB script based on Fieldtrip toolbox^35^. First, contacts located in the seizure onset zone or those with evident epileptiform artifacts were excluded, with the process carefully reviewed and confirmed by experienced clinicians. For each of the remaining contacts, signals during the TMS artifact period [-10 ms, 25 ms] were removed, resulting in segments of iEEG signal ranging from 26 ms to 1990 ms post-TMS pulse onset.

To better control the decay artifacts associated with TMS, which last approximately 150 ms post-TMS stimulus onset, we conducted the filtering process to each segmented iEEG signal rather than the original intact signals contaminated by TMS artifacts. Filtering the intact signals would introduce enhanced drifts during the baseline period preceding the TMS onset. The filters used included a notch filter (3^rd^ order Butterworth filter, cutoff frequencies 57-63 Hz and its seven harmonics) and a bandpass filter (2^nd^ order Butterworth IIR filter, 6 dB cutoff frequency at 2 Hz and 35 Hz).

Following the filtering process, signals during the TMS artifact period were interpolated with filtered stationary iEEG signals that represented similar pattern as the background signal^36^. Specifically, signals with equal length to the artifact period immediately preceding and following the artifact period were first extracted and reversed. A tapering matrix (1:1/n:0 for the preceding data, 0:1/n:1 for the following data, where n is the number of samples in the artifact period) was further applied to the reversed signals. The two resultant signals were subsequently summed together and used to replace the artifact period. After interpolation, the iEEG signals were downsampled to 1000 Hz, clipped into epochs from -200ms to 500ms, and baseline-corrected by subtracting the pre-stimulus voltage average between –200 ms and –50 ms. Next, we addressed the remaining decay artifact by applying the Adaptive Detrend Algorithm (ADA), as outlined in earlier research^37^ and done in our previous work^19^.

After stimulation artifact removal, trials that contained non-physiological signals (e.g., cable motion artifact, long decay artifact from amplifier saturation) or potential interictal spikes were discarded. The criteria for the latter were defined as those exceeding 10 times the baseline standard deviations (SDs) or 100 µV at 25 ms (i.e., the end of TMS artifact), or those exceeding 50 times the baseline SDs or 200 µV throughout the entire epoch.

### Quantification and Significance Testing of intracranial TMS-Evoked Potentials (iTEPs)

We determined the iTEP significance of a contact if it met the following two criteria: first, its response amplitude exceeded 5 SDs with respect to the baseline following active TMS but not following sham TMS. Second, its response following active TMS was significantly different from that following sham TMS through a nonparametric clustering method^19,35^.

For the first criterion, we averaged preprocessed epochs across all the remaining trials (46-50 trials based on trials discarded) for each contact. The resultant iTEPs were then z-scored with respect to the baseline. For the second criterion, we compared active and sham iTEPs at each time point of [25 ms, 500 ms] post-TMS to identify clusters of significant time points with paired *t-*tests at an alpha level of 0.05. A cluster was defined as a period with a contiguous sequence of significant time points (cluster threshold > 10 ms). The “total statistic” for each cluster was determined by summing the *t*-statistics within the cluster. A contact’s final cluster statistic is defined as the sum of the largest three clusters’ “total statistic.” A null distribution was further generated by computing cluster statistics for randomly shuffled trials (assigned to either the active or sham TMS condition) with 1000 simulations. The actual cluster statistic of each contact was finally compared with this null distribution and a contact’s iTEP was considered significant using *p* < 0.05 (after FDR correction for multiple contacts comparison).

### Corticocortical evoked potential stimulation and analysis

The corticocortical evoked potential (CCEP) experiment for subject 460 delivered 60 biphasic pulses of bipolar electrical stimulation (4.5 mA, 200 µs pulse width, jittered around 0.5 Hz) through two neighboring electrode contacts on a surface electrocorticography grid over the patient’s left prefrontal cortex (see **Supplementary Fig. S3**). The recording parameters were the same as the TMS-iEEG experiments.

iEEG preprocessing steps for the CCEP analysis followed the same steps as the iTEP preprocessing, except with the following notable differences: 1) shorter [-8 ms, 8 ms] iES artifact period during which signals were removed and interpolated with synthesized stationary iEEG signals; 2) consequently, wider time window for baseline correction [-200 ms, -10 ms] due to the shorter artifact; 3) filtering applied to the entire iEEG signal of each contact, as opposed to segment-based filtering; 4) no detrending step required.

After preprocessing, contact average EP were calculated by averaging across the remaining trials for each electrode and then z-score normalized with respect to their baseline [-200ms, -10ms] SD. To elucidate whether the iES at DLPFC evoked significant responses in the sgACC contact, we applied one-sample t-test at each post-stimulation timepoint within [9 ms, 500 ms] and then FDR corrected (*p* < 0.05) across all these timepoints to determine the significant timepoints.

### Functional connectivity assessment

We calculated functional connectivity (FC) between the DLPFC TMS sites and sgACC contact sites using preprocessed resting-state functional MRI data. First, we created a 4 mm radius spherical mask centered on MNI coordinates of each TMS site and electrode contact, excluding voxels located outside the brain. For the masks of TMS sites, we projected the scalp stimulation site to the nearest vertex on the pial surface using Brainsight data to inform the average stimulation trajectory. Next, we averaged the BOLD signal within each mask to obtain one time course of BOLD signal for this mask. The TMS-to-contact FC was defined as the Pearson correlation between the BOLD time course of each contact and the corresponding TMS site.

## Acknowledgements

We would like to acknowledge the patients and families who graciously agreed to participate in this research, as well as research members in the Human Brain Research Laboratory who assisted with data collection and provided feedback on the project including Ariane Rhone, Haiming Chen, Christopher Kovach, Eric Tsang, Ben Pace, Brandt Uitermarkt, Phil Gander, Joel Berger, Hiroyuki Oya, Chris Garcia, and Matthew Howard.

