## Supplemental Table S1 for "Dorsolateral prefrontal cortex TMS evokes responses in the subgenual anterior cingulate cortex: Evidence from human intracranial EEG"

**Table S1. Significance results of whole-brain electrode contacts for Patient 460**

| Contact | MNI-x | MNI-y | MNI-z | AAL_label | Harvard-Oxford_cortical+subcortical_label | TMS_peak (SD) | Sham_peak (SD) | T (3 largest clusters) | P (clusters) | Significance* (1-Yes, 0-No) |
|---|---|---|---|---|---|---|---|---|---|---|
| 1 | -11.6 | 41.2 | -27.3 | OFCmed_L | Frontal_Pole | 4.33 | 4.09 | 5.99 | 0.6620 | 0 |
| 2 | -22.5 | 38.0 | -21.1 | OFCant_L | Frontal_Pole | 7.16 | 3.72 | 78.31 | 0.2850 | 0 |
| 3 | -36.2 | 39.4 | -18.5 | OFCant_L | Frontal_Pole | 8.30 | 0.95 | 170.91 | 0.0660 | 0 |
| 4 | -44.7 | 38.6 | -19.9 | None | Frontal_Pole | 27.18 | 1.46 | 75.24 | 0.2800 | 0 |
| 21 | 31.8 | 30.0 | -22.4 | OFCpost_R | Frontal_Orbital_Cortex | 4.84 | 3.77 | 80.99 | 0.2310 | 0 |
| 22 | 21.4 | 30.0 | -23.6 | OFCmed_R | Frontal_Orbital_Cortex | 4.16 | 2.80 | 88.45 | 0.2140 | 0 |
| 23 | 10.6 | 29.3 | -27.4 | OFCmed_R | Frontal_Orbital_Cortex | 4.02 | 3.79 | 160.41 | 0.0340 | 0 |
| 24 | 1.5 | 29.3 | -28.1 | Rectus_L | Subcallosal_Cortex | 4.52 | 4.55 | 54.26 | 0.3650 | 0 |
| 25 | -11.4 | 29.1 | -27.3 | OFCmed_L | Frontal_Orbital_Cortex | 1.91 | 5.94 | 229.10 | 0.0430 | 0 |
| 26 | -22.6 | 29.6 | -23.2 | None | Frontal_Orbital_Cortex | 2.72 | 4.09 | 193.09 | 0.0450 | 0 |
| 27 | -34.6 | 27.4 | -21.8 | None | Frontal_Orbital_Cortex | 3.27 | 2.12 | 97.20 | 0.1980 | 0 |
| 28 | -43.5 | 27.5 | -16.9 | OFClat_L | Frontal_Orbital_Cortex | 2.92 | 2.09 | 94.46 | 0.2280 | 0 |
| 33 | -6.1 | 21.9 | -16.5 | Rectus_L | Subcallosal_Cortex | 3.12 | 1.50 | 174.67 | 0.0880 | 0 |
| 34 | -8.2 | 26.8 | -8.2 | None | Subcallosal_Cortex | 5.20 | 1.13 | 276.84 | 0.0280 | 1 |
| 37 | -12.1 | 36.4 | 25.0 | Frontal_Sup_Medial_L | Paracingulate_Gyrus | 7.39 | 1.10 | 232.75 | 0.0420 | 1 |
| 38 | -13.5 | 41.6 | 37.8 | Frontal_Sup_2_L | Frontal_Pole | 4.26 | 0.96 | 18.09 | 0.5730 | 0 |
| 39 | -13.5 | 46.0 | 48.7 | Frontal_Sup_2_L | Frontal_Pole | 9.27 | 1.67 | 120.41 | 0.1590 | 0 |
| 41 | -9.3 | 21.3 | 25.6 | None | Cingulate_Gyrus,_anterior_division | 7.63 | 1.06 | 93.25 | 0.2110 | 0 |
| 42 | -10.9 | 22.5 | 29.5 | None | Paracingulate_Gyrus | 6.72 | 2.10 | 188.60 | 0.0490 | 1 |
| 46 | -13.8 | 32.7 | 50.0 | Frontal_Sup_2_L | Superior_Frontal_Gyrus | 2.87 | 2.21 | 112.13 | 0.1660 | 0 |
| 47 | -14.5 | 34.1 | 54.9 | Frontal_Sup_2_L | Superior_Frontal_Gyrus | 3.40 | 1.84 | 79.10 | 0.2730 | 0 |
| 50 | -37.6 | -22.4 | 1.2 | None | Insular_Cortex | 23.14 | 17.51 | 413.17 | 0.0000 | 0 |
| 51 | -42.6 | -22.2 | 1.5 | Temporal_Sup_L | Heschls_Gyrus_(includes_H1_and_H2) | 17.87 | 15.40 | 387.40 | 0.0000 | 0 |
| 52 | -47.6 | -21.9 | 2.4 | Temporal_Sup_L | Heschls_Gyrus_(includes_H1_and_H2) | 9.08 | 6.42 | 359.00 | 0.0000 | 0 |
| 53 | -52.9 | -21.7 | 3.7 | Temporal_Sup_L | Planum_Temporale | 9.87 | 4.12 | 305.15 | 0.0010 | 1 |
| 54 | -57.7 | -21.1 | 5.0 | Temporal_Sup_L | Planum_Temporale | 6.03 | 2.66 | 210.39 | 0.0150 | 1 |
| 56 | -67.7 | -20.0 | 6.2 | Temporal_Sup_L | Superior_Temporal_Gyrus,_posterior_division | 6.54 | 4.55 | 215.84 | 0.0190 | 1 |

| | | | | | | | | | | |
|---|---|---|---|---|---|---|---|---|---|---|
| 57 | -23.0 | -56.7 | -16.4 | Fusiform_L | Temporal_Occipital_Fusiform_Cortex | 1.59 | 2.33 | 69.90 | 0.3200 | 0 |
| 60 | -51.4 | -41.8 | -27.0 | Temporal_Inf_L | Inferior_Temporal_Gyrus,_posterior_division | 7.33 | 2.48 | 247.74 | 0.0130 | 1 |
| 65 | -2.7 | 10.4 | 31.7 | Cingulate_Mid_L | Cingulate_Gyrus,_anterior_division | 14.52 | 2.66 | 617.65 | 0.0000 | 1 |
| 66 | -5.6 | 6.9 | 43.8 | Supp_Motor_Area_L | Paracingulate_Gyrus | 5.60 | 4.92 | 48.44 | 0.3940 | 0 |
| 67 | -9.0 | 7.2 | 53.7 | Supp_Motor_Area_L | Juxtapositional_Lobule_Cortex_(formerly_Supplementary_Motor_Cortex) | 9.96 | 3.89 | 382.77 | 0.0010 | 1 |
| 68 | -14.6 | 4.1 | 62.6 | Supp_Motor_Area_L | Superior_Frontal_Gyrus | 11.08 | 3.13 | 181.74 | 0.0390 | 1 |
| 70 | -36.0 | -25.2 | 14.2 | Heschl_L | Heschls_Gyrus_(includes_H1_and_H2) | 20.63 | 16.73 | 601.18 | 0.0000 | 0 |
| 71 | -41.9 | -17.0 | 7.7 | Heschl_L | Heschls_Gyrus_(includes_H1_and_H2) | 11.09 | 4.45 | 658.78 | 0.0000 | 1 |
| 72 | -47.9 | -11.1 | 3.0 | Temporal_Sup_L | Heschls_Gyrus_(includes_H1_and_H2) | 18.01 | 3.83 | 294.11 | 0.0020 | 1 |
| 74 | -31.0 | 15.6 | 13.7 | None | Frontal_Operculum_Cortex | 8.82 | 1.71 | 355.65 | 0.0030 | 1 |
| 81 | -15.2 | -44.9 | -4.0 | Lingual_L | Lingual_Gyrus | 1.25 | 2.72 | 0.00 | 0.6950 | 0 |
| 84 | -26.4 | -34.7 | -8.8 | Hippocampus_L | Left_Hippocampus | 9.14 | 3.75 | 260.10 | 0.0050 | 1 |
| 85 | -30.2 | -30.6 | -10.3 | Hippocampus_L | Left_Hippocampus | 8.30 | 4.09 | 240.36 | 0.0240 | 1 |
| 86 | -34.0 | -27.0 | -11.8 | Hippocampus_L | Left_Hippocampus | 4.66 | 3.55 | 385.64 | 0.0020 | 0 |
| 97 | -30.2 | 8.1 | -18.4 | Insula_L | Frontal_Orbital_Cortex | 4.47 | 1.45 | 328.02 | 0.0030 | 0 |
| 98 | -40.5 | 9.5 | -15.7 | Temporal_Pole_Sup_L | nan | 6.76 | 3.02 | 302.94 | 0.0040 | 1 |
| 99 | -51.8 | 9.6 | -15.1 | Temporal_Pole_Sup_L | Temporal_Pole | 3.37 | 3.93 | 0.00 | 0.5520 | 0 |
| 101 | -14.0 | -9.5 | -19.4 | Hippocampus_L | Left_Hippocampus | 2.79 | 4.25 | 17.95 | 0.5590 | 0 |
| 102 | -16.5 | -8.8 | -19.7 | Hippocampus_L | Left_Hippocampus | 3.33 | 3.53 | 40.91 | 0.4450 | 0 |
| 103 | -19.3 | -8.1 | -20.2 | Hippocampus_L | Left_Hippocampus | 4.37 | 3.53 | 35.30 | 0.4620 | 0 |
| 104 | -21.9 | -7.5 | -20.8 | Hippocampus_L | Left_Amygdala | 11.88 | 3.53 | 43.07 | 0.3870 | 0 |
| 105 | -24.4 | -6.8 | -21.4 | Hippocampus_L | Left_Amygdala | 10.25 | 3.98 | 0.00 | 0.6160 | 0 |
| 106 | -27.0 | -6.1 | -21.9 | Amygdala_L | Left_Amygdala | 4.59 | 7.09 | 0.00 | 0.5900 | 0 |
| 107 | -29.5 | -5.4 | -22.3 | Amygdala_L | Left_Amygdala | 3.26 | 7.51 | 0.00 | 0.5460 | 0 |
| 109 | -25.0 | 25.1 | -20.4 | OFCpost_L | Frontal_Orbital_Cortex | 4.00 | 1.73 | 327.48 | 0.0050 | 0 |
| 113 | -32.8 | 47.4 | 16.6 | Frontal_Mid_2_L | Frontal_Pole | 15.07 | 3.23 | 91.56 | 0.2370 | 0 |

| | | | | | | | | | | |
|---|---|---|---|---|---|---|---|---|---|---|
| 114 | -34.5 | 50.5 | 27.3 | Frontal_Mid_2_L | Frontal_Pole | 6.13 | 1.64 | 49.26 | 0.3720 | 0 |
| 131 | -43.2 | 5.2 | -46.3 | None | Temporal_Pole | 1.58 | 2.02 | 148.98 | 0.0950 | 0 |
| 132 | -47.2 | 12.7 | -41.1 | None | Temporal_Pole | 3.96 | 3.93 | 134.08 | 0.1210 | 0 |
| 133 | -50.4 | 14.9 | -31.2 | Temporal_Pole_Mid_L | Temporal_Pole | 3.22 | 3.10 | 186.98 | 0.0480 | 0 |
| 134 | -51.8 | 17.8 | -22.7 | None | Temporal_Pole | 7.71 | 2.50 | 82.15 | 0.2370 | 0 |
| 137 | -47.8 | -2.2 | -46.5 | None | Inferior_Temporal_Gyrus,_anterior_division | 2.06 | 2.06 | 53.45 | 0.4110 | 0 |
| 138 | -53.3 | 3.9 | -39.0 | None | Temporal_Pole | 2.43 | 2.28 | 0.00 | 0.7560 | 0 |
| 139 | -56.3 | 8.5 | -30.2 | Temporal_Mid_L | Temporal_Pole | 5.50 | 2.91 | 87.70 | 0.2630 | 0 |
| 140 | -34.9 | -21.0 | -34.0 | None | Temporal_Fusiform_Cortex,_posterior_division | 6.60 | 1.59 | 358.17 | 0.0030 | 1 |
| 141 | -44.0 | -16.8 | -37.7 | None | Inferior_Temporal_Gyrus,_posterior_division | 3.36 | 5.54 | 154.97 | 0.0670 | 0 |
| 143 | -57.6 | -7.0 | -36.5 | Temporal_Inf_L | Inferior_Temporal_Gyrus,_anterior_division | 6.55 | 3.88 | 0.00 | 0.7180 | 0 |
| 144 | -37.0 | -25.3 | -31.1 | None | Temporal_Fusiform_Cortex,_posterior_division | 1.78 | 2.62 | 90.51 | 0.2430 | 0 |
| 145 | -46.8 | -21.2 | -33.6 | Temporal_Inf_L | Inferior_Temporal_Gyrus,_posterior_division | 4.25 | 4.63 | 193.18 | 0.0300 | 0 |
| 148 | -43.1 | -36.0 | -27.8 | Temporal_Inf_L | Temporal_Fusiform_Cortex,_posterior_division | 9.60 | 2.46 | 128.44 | 0.1330 | 0 |
| 149 | -52.8 | -31.3 | -28.9 | Temporal_Inf_L | Inferior_Temporal_Gyrus,_posterior_division | 3.67 | 4.09 | 188.33 | 0.0340 | 0 |
| 150 | -60.3 | -26.1 | -30.4 | None | Inferior_Temporal_Gyrus,_posterior_division | 3.01 | 2.81 | 258.38 | 0.0060 | 0 |
| 151 | -64.2 | -18.4 | -28.8 | Temporal_Inf_L | nan | 4.87 | 1.95 | 97.17 | 0.2300 | 0 |
| 161 | -9.3 | 68.2 | 21.8 | None | Frontal_Pole | 3.44 | 1.71 | 159.92 | 0.0940 | 0 |
| 162 | -11.0 | 62.8 | 30.1 | Frontal_Sup_Medial_L | Frontal_Pole | 2.58 | 2.10 | 0.00 | 0.5930 | 0 |
| 163 | -15.9 | 55.7 | 38.3 | Frontal_Sup_2_L | Frontal_Pole | 2.95 | 1.25 | 126.99 | 0.1550 | 0 |
| 164 | -18.6 | 46.1 | 46.7 | Frontal_Sup_2_L | Frontal_Pole | 9.94 | 1.38 | 124.20 | 0.1650 | 0 |
| 165 | -19.4 | 36.5 | 55.1 | Frontal_Sup_2_L | nan | 3.95 | 1.71 | 56.08 | 0.3710 | 0 |
| 166 | -21.2 | 25.6 | 61.6 | Frontal_Sup_2_L | Superior_Frontal_Gyrus | 2.50 | 2.94 | 0.00 | 0.7280 | 0 |
| 167 | -21.3 | 14.0 | 67.9 | Frontal_Sup_2_L | Superior_Frontal_Gyrus | 5.42 | 2.59 | 90.72 | 0.2320 | 0 |
| 168 | -22.8 | 1.8 | 72.4 | Frontal_Sup_2_L | Superior_Frontal_Gyrus | 5.85 | 2.14 | 59.01 | 0.3750 | 0 |

| | | | | | | | | | | |
|---|---|---|---|---|---|---|---|---|---|---|
| 169 | -21.0 | 66.3 | 18.2 | Frontal_Sup_2_L | Frontal_Pole | 1.96 | 1.40 | 49.51 | 0.3340 | 0 |
| 170 | -22.0 | 63.3 | 25.1 | Frontal_Sup_2_L | Frontal_Pole | 2.34 | 1.79 | 0.00 | 0.6120 | 0 |
| 171 | -24.5 | 55.3 | 31.8 | Frontal_Sup_2_L | Frontal_Pole | 8.32 | 3.20 | 314.88 | 0.0050 | 1 |
| 172 | -28.1 | 43.6 | 43.3 | Frontal_Sup_2_L | nan | 11.00 | 2.72 | 240.13 | 0.0130 | 1 |
| 173 | -29.8 | 35.0 | 48.9 | Frontal_Mid_2_L | nan | 3.99 | 3.42 | 186.10 | 0.0310 | 0 |
| 175 | -32.1 | 13.0 | 63.1 | Frontal_Mid_2_L | Middle_Frontal_Gyrus | 3.69 | 2.68 | 214.34 | 0.0190 | 0 |
| 176 | -33.0 | 2.2 | 65.3 | Frontal_Mid_2_L | Middle_Frontal_Gyrus | 2.54 | 3.98 | 265.69 | 0.0030 | 0 |
| 193 | -26.5 | 67.1 | 7.9 | Frontal_Sup_2_L | Frontal_Pole | 2.18 | 3.89 | 0.00 | 0.5670 | 0 |
| 194 | -34.0 | 60.3 | 16.1 | Frontal_Mid_2_L | Frontal_Pole | 2.84 | 6.38 | 0.00 | 0.6320 | 0 |
| 195 | -35.8 | 55.0 | 24.4 | Frontal_Mid_2_L | Frontal_Pole | 4.61 | 7.97 | 373.25 | 0.0030 | 0 |
| 196 | -38.5 | 46.1 | 31.9 | Frontal_Mid_2_L | Frontal_Pole | 10.89 | 2.59 | 423.85 | 0.0000 | 1 |
| 197 | -40.0 | 37.1 | 38.8 | Frontal_Mid_2_L | nan | 5.51 | 2.55 | 111.56 | 0.1560 | 0 |
| 198 | -42.5 | 24.9 | 47.6 | None | Middle_Frontal_Gyrus | 4.36 | 2.24 | 204.78 | 0.0230 | 0 |
| 199 | -42.2 | 15.3 | 55.5 | Frontal_Mid_2_L | Middle_Frontal_Gyrus | 3.55 | 1.93 | 152.41 | 0.0590 | 0 |
| 200 | -40.2 | 4.3 | 60.2 | Frontal_Mid_2_L | Middle_Frontal_Gyrus | 2.08 | 2.11 | 108.71 | 0.1560 | 0 |
| 201 | -35.9 | 63.0 | 1.7 | None | Frontal_Pole | 7.32 | 3.83 | 47.19 | 0.4030 | 0 |
| 202 | -39.4 | 59.2 | 9.5 | Frontal_Mid_2_L | Frontal_Pole | 4.65 | 3.73 | 66.37 | 0.3220 | 0 |
| 203 | -42.9 | 51.4 | 17.4 | Frontal_Mid_2_L | Frontal_Pole | 4.37 | 10.52 | 307.00 | 0.0030 | 0 |
| 204 | -46.4 | 42.4 | 24.1 | None | Frontal_Pole | 6.50 | 2.99 | 174.21 | 0.0590 | 0 |
| 205 | -49.1 | 32.5 | 31.9 | None | Middle_Frontal_Gyrus | 3.99 | 2.12 | 16.32 | 0.6470 | 0 |
| 206 | -50.1 | 21.1 | 41.7 | Frontal_Mid_2_L | Middle_Frontal_Gyrus | 4.35 | 1.84 | 59.00 | 0.3770 | 0 |
| 207 | -48.2 | 10.8 | 48.6 | Frontal_Mid_2_L | Middle_Frontal_Gyrus | 3.67 | 2.56 | 227.50 | 0.0040 | 0 |
| 208 | -49.1 | 0.6 | 54.1 | None | Precentral_Gyrus | 7.67 | 4.95 | 189.62 | 0.0280 | 1 |
| 209 | -41.3 | 58.5 | -5.7 | Frontal_Mid_2_L | Frontal_Pole | 6.27 | 3.56 | 87.17 | 0.2370 | 0 |
| 210 | -45.2 | 53.6 | -0.1 | Frontal_Mid_2_L | Frontal_Pole | 2.86 | 4.21 | 144.99 | 0.1060 | 0 |
| 211 | -49.0 | 45.8 | 9.2 | Frontal_Inf_Tri_L | Frontal_Pole | 2.26 | 2.84 | 60.18 | 0.3500 | 0 |
| 213 | -54.6 | 27.1 | 24.6 | Frontal_Inf_Tri_L | nan | 3.41 | 2.61 | 170.19 | 0.0630 | 0 |
| 214 | -53.8 | 18.0 | 34.8 | None | Middle_Frontal_Gyrus | 2.92 | 1.96 | 23.22 | 0.5900 | 0 |
| 215 | -54.5 | 7.0 | 40.4 | Precentral_L | Precentral_Gyrus | 4.23 | 3.95 | 10.28 | 0.7360 | 0 |
| 216 | -55.5 | -1.6 | 47.8 | Precentral_L | nan | 3.50 | 7.07 | 81.59 | 0.2630 | 0 |
| 217 | -45.0 | 52.0 | -11.2 | None | Frontal_Pole | 3.27 | 4.63 | 118.54 | 0.1700 | 0 |
| 218 | -49.9 | 46.2 | -6.1 | None | Frontal_Pole | 2.75 | 2.86 | 20.33 | 0.6450 | 0 |
| 219 | -54.3 | 38.3 | 1.0 | None | Frontal_Pole | 1.91 | 1.89 | 64.09 | 0.3540 | 0 |

| | | | | | | | | | | |
|---|---|---|---|---|---|---|---|---|---|---|
| 220 | -56.6 | 30.4 | 9.7 | None | Inferior_Frontal_Gyrus,_pars_triangularis | 4.03 | 5.97 | 0.00 | 0.7500 | 0 |
| 222 | -60.1 | 10.6 | 26.5 | None | Precentral_Gyrus | 4.84 | 2.07 | 40.51 | 0.5010 | 0 |
| 223 | -60.8 | 4.0 | 33.9 | None | Precentral_Gyrus | 4.60 | 2.62 | 192.48 | 0.0070 | 0 |
| 224 | -61.1 | -7.2 | 39.5 | Postcentral_L | nan | 4.37 | 3.19 | 101.05 | 0.1270 | 0 |
| 225 | -62.0 | 7.3 | 8.4 | None | Precentral_Gyrus | 15.58 | 3.31 | 481.61 | 0.0000 | 1 |
| 226 | -63.9 | -1.2 | 14.3 | None | Precentral_Gyrus | 10.93 | 4.24 | 667.28 | 0.0000 | 1 |
| 227 | -67.0 | -9.8 | 20.4 | None | Postcentral_Gyrus | 4.11 | 2.55 | 158.20 | 0.0300 | 0 |
| 228 | -67.2 | -21.1 | 25.4 | None | Postcentral_Gyrus | 2.42 | 2.66 | 282.37 | 0.0000 | 0 |
| 229 | -68.0 | -30.1 | 29.5 | SupraMarginal_L | nan | 4.82 | 2.34 | 0.00 | 0.7790 | 0 |
| 230 | -66.3 | -41.2 | 34.8 | SupraMarginal_L | nan | 6.25 | 2.96 | 47.14 | 0.4940 | 0 |
| 235 | -67.0 | -17.3 | 8.9 | Temporal_Sup_L | nan | 7.30 | 4.00 | 369.15 | 0.0000 | 1 |
| 238 | -66.5 | -45.1 | 25.4 | SupraMarginal_L | Supramarginal_Gyrus,_posterior_division | 9.69 | 3.38 | 88.19 | 0.2640 | 0 |
| 241 | -63.2 | -2.3 | -12.0 | Temporal_Mid_L | Middle_Temporal_Gyrus,_anterior_division | 11.99 | 2.16 | 213.85 | 0.0360 | 1 |
| 242 | -65.6 | -10.9 | -4.0 | Temporal_Mid_L | nan | 5.62 | 2.08 | 307.28 | 0.0050 | 1 |
| 243 | -68.4 | -20.7 | -0.2 | Temporal_Mid_L | Superior_Temporal_Gyrus,_posterior_division | 5.73 | 2.45 | 193.12 | 0.0310 | 1 |
| 244 | -69.2 | -30.9 | 4.8 | Temporal_Mid_L | Superior_Temporal_Gyrus,_posterior_division | 3.98 | 7.74 | 65.83 | 0.3430 | 0 |
| 247 | -62.4 | -58.6 | 20.2 | Temporal_Mid_L | Angular_Gyrus | 5.52 | 1.70 | 148.33 | 0.0770 | 0 |
| 248 | -57.7 | -67.5 | 24.7 | None | Lateral_Occipital_Cortex,_superior_division | 4.79 | 2.82 | 188.05 | 0.0260 | 0 |
| 249 | -65.1 | -4.9 | -19.5 | None | Middle_Temporal_Gyrus,_anterior_division | 4.47 | 2.50 | 308.43 | 0.0020 | 0 |
| 250 | -68.7 | -15.2 | -16.1 | None | Middle_Temporal_Gyrus,_posterior_division | 4.53 | 1.50 | 26.33 | 0.6000 | 0 |
| 251 | -69.5 | -24.8 | -11.7 | None | Middle_Temporal_Gyrus,_posterior_division | 4.53 | 1.79 | 0.00 | 0.7470 | 0 |
| 254 | -66.3 | -53.7 | 4.2 | None | Middle_Temporal_Gyrus,_temporooccipital_part | 6.59 | 2.44 | 218.71 | 0.0140 | 1 |

***:** If active TMS response > ± 5 SDs baseline, sham TMS response < ± 5 SDs baseline, and active TMS response was significantly stronger in nonparametric clustering analysis with 1000 permutations, then significance = 1 otherwise 0
