## Supplemental Table S2 for "Dorsolateral prefrontal cortex TMS evokes responses in the subgenual anterior cingulate cortex: Evidence from human intracranial EEG"

**Table S2. Significance results of whole-brain electrode contacts for Patient 625**

| Contact | MNI-x | MNI-y | MNI-z | AAL_label | Harvard-Oxford_cortical+subcortical_label | TMS_peak (SD) | Sham_peak (SD) | T (3 largest clusters) | P (clusters) | Significance* (1-Yes, 0-No) |
|---|---|---|---|---|---|---|---|---|---|---|
| 1 | -7.1 | 15.2 | -14.2 | Rectus_L | Subcallosal_Cortex | 9.60 | 3.95 | 886.16 | 0.0000 | 1 |
| 7 | -29.4 | 25.3 | -6.3 | Insula_L | Frontal_Orbital_Cortex | 8.73 | 4.52 | 652.81 | 0.0000 | 1 |
| 8 | -35.7 | 27.7 | -6.0 | Frontal_Inf_Orb_2_L | Frontal_Orbital_Cortex | 13.29 | 3.42 | 636.06 | 0.0000 | 1 |
| 9 | -42.2 | 32.6 | -5.9 | Frontal_Inf_Orb_2_L | Frontal_Orbital_Cortex | 34.11 | 4.20 | 810.74 | 0.0000 | 1 |
| 10 | -48.7 | 34.6 | -5.4 | Frontal_Inf_Orb_2_L | Frontal_Pole | 14.95 | 10.93 | 1094.14 | 0.0000 | 0 |
| 11 | -53.4 | 35.0 | -4.1 | None | Inferior_Frontal_Gyrus,_pars_triangularis | 20.87 | 5.07 | 665.22 | 0.0000 | 0 |
| 16 | -26.3 | 52.8 | 6.1 | Frontal_Mid_2_L | Frontal_Pole | 13.75 | 2.00 | 822.78 | 0.0000 | 1 |
| 17 | -32.1 | 54.3 | 7.6 | Frontal_Mid_2_L | Frontal_Pole | 13.57 | 2.26 | 702.21 | 0.0000 | 1 |
| 18 | -38.0 | 55.4 | 8.4 | Frontal_Mid_2_L | Frontal_Pole | 8.75 | 3.76 | 606.73 | 0.0000 | 1 |
| 19 | -43.2 | 55.6 | 9.2 | None | Frontal_Pole | 18.13 | 3.59 | 747.89 | 0.0000 | 1 |
| 23 | -4.8 | 34.9 | 29.4 | Frontal_Sup_Medial_L | Paracingulate_Gyrus | 20.48 | 2.67 | 1161.52 | 0.0000 | 1 |
| 24 | -9.7 | 35.8 | 30.0 | Frontal_Sup_Medial_L | Paracingulate_Gyrus | 20.29 | 2.38 | 1381.31 | 0.0000 | 1 |
| 26 | -23.0 | 35.5 | 30.4 | Frontal_Sup_2_L | Middle_Frontal_Gyrus | 10.79 | 2.34 | 1489.93 | 0.0000 | 1 |
| 27 | -31.9 | 37.0 | 31.6 | Frontal_Mid_2_L | Middle_Frontal_Gyrus | 14.79 | 3.30 | 1354.51 | 0.0000 | 1 |
| 28 | -39.2 | 39.0 | 31.7 | Frontal_Mid_2_L | Frontal_Pole | 11.02 | 4.12 | 1316.27 | 0.0000 | 1 |
| 33 | -5.3 | -46.6 | 25.4 | Cingulate_Post_L | Cingulate_Gyrus,_posterior_division | 11.30 | 4.12 | 213.28 | 0.0290 | 1 |
| 41 | -54.1 | -48.6 | 30.0 | SupraMarginal_L | Supramarginal_Gyrus,_posterior_division | 11.54 | 3.86 | 617.91 | 0.0000 | 1 |
| 42 | -59.4 | -48.8 | 28.3 | SupraMarginal_L | Supramarginal_Gyrus,_posterior_division | 9.97 | 4.35 | 584.58 | 0.0000 | 1 |
| 45 | -8.5 | -19.2 | 59.1 | Supp_Motor_Area_L | Precentral_Gyrus | 33.97 | 9.40 | 1379.56 | 0.0000 | 0 |
| 47 | -22.3 | -15.1 | 56.7 | Precentral_L | Precentral_Gyrus | 29.38 | 4.71 | 1418.48 | 0.0000 | 1 |
| 48 | -28.0 | -11.9 | 55.7 | Precentral_L | Precentral_Gyrus | 21.01 | 3.98 | 1165.07 | 0.0000 | 1 |
| 49 | -34.9 | -11.1 | 55.2 | Precentral_L | Precentral_Gyrus | 14.76 | 4.33 | 1053.85 | 0.0000 | 1 |
| 50 | -42.0 | -11.1 | 56.2 | Precentral_L | Precentral_Gyrus | 45.85 | 4.50 | 1306.61 | 0.0000 | 1 |
| 51 | -47.1 | -9.6 | 57.1 | Precentral_L | Precentral_Gyrus | 54.61 | 5.57 | 1215.24 | 0.0000 | 0 |
| 52 | -50.4 | -6.8 | 56.4 | None | Precentral_Gyrus | 30.58 | 9.26 | 1221.24 | 0.0000 | 0 |

| | | | | | | | | | | |
|---|---|---|---|---|---|---|---|---|---|---|
| 55 | -2.6 | -9.3 | 34.5 | Cingulate_Mid_L | Cingulate_Gyrus,_anterior_division | 12.98 | 4.86 | 1254.86 | 0.0000 | 1 |
| 61 | -40.4 | 3.0 | 33.4 | Precentral_L | Precentral_Gyrus | 27.09 | 9.33 | 1365.05 | 0.0000 | 0 |
| 62 | -46.2 | 6.1 | 32.5 | Precentral_L | Middle_Frontal_Gyrus | 42.42 | 9.15 | 2014.40 | 0.0000 | 0 |
| 64 | -57.7 | 7.3 | 35.1 | Precentral_L | Precentral_Gyrus | 18.79 | 5.70 | 1418.03 | 0.0000 | 0 |
| 75 | -38.4 | 3.4 | -17.1 | None | Insular_Cortex | 8.87 | 9.42 | 827.30 | 0.0000 | 0 |
| 76 | -37.9 | 6.9 | -10.5 | Insula_L | Insular_Cortex | 21.85 | 5.12 | 1287.00 | 0.0000 | 0 |
| 77 | -37.2 | 10.4 | -5.9 | Insula_L | Insular_Cortex | 26.59 | 5.74 | 1309.68 | 0.0000 | 0 |
| 78 | -36.9 | 14.1 | -2.1 | Insula_L | Insular_Cortex | 23.51 | 6.70 | 1258.82 | 0.0000 | 0 |
| 79 | -37.1 | 16.7 | 2.3 | Insula_L | Insular_Cortex | 14.34 | 2.62 | 598.32 | 0.0000 | 1 |
| 80 | -38.0 | 19.5 | 6.8 | Frontal_Inf_Tri_L | Frontal_Operculum_Cortex | 16.37 | 3.37 | 592.22 | 0.0000 | 1 |
| 83 | -41.7 | 27.6 | 21.6 | Frontal_Inf_Tri_L | Middle_Frontal_Gyrus | 3.99 | 3.92 | 550.80 | 0.0000 | 0 |
| 84 | -40.6 | 29.5 | 27.1 | Frontal_Mid_2_L | Middle_Frontal_Gyrus | 14.84 | 3.38 | 1755.84 | 0.0000 | 1 |
| 85 | -36.9 | 31.1 | 34.4 | Frontal_Mid_2_L | Middle_Frontal_Gyrus | 14.71 | 3.09 | 1450.07 | 0.0000 | 1 |
| 86 | -32.7 | 35.5 | 41.6 | Frontal_Mid_2_L | Middle_Frontal_Gyrus | 13.09 | 2.66 | 926.98 | 0.0000 | 1 |
| 87 | -34.5 | 13.9 | -9.5 | Insula_L | Insular_Cortex | 23.15 | 4.49 | 1302.51 | 0.0000 | 1 |
| 88 | -41.1 | 14.6 | -11.2 | Insula_L | Insular_Cortex | 20.48 | 5.34 | 969.03 | 0.0000 | 0 |
| 89 | -46.9 | 14.9 | -13.7 | Temporal_Pole_Sup_L | Temporal_Pole | 7.66 | 3.06 | 351.76 | 0.0030 | 1 |
| 90 | -52.0 | 16.0 | -15.1 | Temporal_Pole_Sup_L | Temporal_Pole | 2.91 | 4.70 | 190.69 | 0.0620 | 0 |
| 91 | -55.7 | 15.2 | -16.7 | None | Temporal_Pole | 5.92 | 8.06 | 340.66 | 0.0120 | 0 |
| 99 | -35.4 | -7.7 | 8.7 | Insula_L | Insular_Cortex | 15.49 | 13.56 | 1461.46 | 0.0000 | 0 |
| 100 | -35.6 | -9.2 | 14.0 | Insula_L | Insular_Cortex | 15.79 | 8.07 | 2126.75 | 0.0000 | 0 |
| 101 | -35.6 | -10.5 | 18.7 | Insula_L | Central_Opercular_Cortex | 21.59 | 28.56 | 1684.19 | 0.0000 | 0 |
| 106 | -33.2 | -18.2 | 47.4 | Precentral_L | Precentral_Gyrus | 28.48 | 6.20 | 869.02 | 0.0000 | 0 |
| 108 | -32.3 | -18.5 | 60.4 | Precentral_L | Precentral_Gyrus | 25.88 | 5.92 | 1044.59 | 0.0000 | 0 |
| 111 | -47.6 | 4.2 | 14.7 | Precentral_L | Precentral_Gyrus | 16.92 | 7.18 | 2665.44 | 0.0000 | 0 |
| 112 | -53.0 | 3.7 | 13.3 | Rolandic_Oper_L | Precentral_Gyrus | 26.27 | 7.21 | 2643.43 | 0.0000 | 0 |
| 113 | -58.8 | 1.9 | 12.6 | Postcentral_L | Precentral_Gyrus | 17.44 | 8.40 | 1183.45 | 0.0000 | 0 |
| 114 | -63.0 | -0.5 | 12.8 | Postcentral_L | Precentral_Gyrus | 33.02 | 7.32 | 2096.70 | 0.0000 | 0 |
| 117 | -32.6 | -24.6 | 9.8 | Heschl_L | Insular_Cortex | 8.98 | 9.17 | 859.68 | 0.0000 | 0 |
| 118 | -38.0 | -23.6 | 8.1 | Heschl_L | Heschls_Gyrus_(includes_H1_and_H2) | 14.09 | 23.45 | 769.57 | 0.0000 | 0 |

| | | | | | | | | | | |
|---|---|---|---|---|---|---|---|---|---|---|
| 119 | -43.6 | -22.1 | 6.9 | Temporal_Sup_L | Heschls_Gyrus_(includes_H1_and_H2) | 25.19 | 21.43 | 433.19 | 0.0000 | 0 |
| 120 | -49.9 | -20.2 | 6.2 | Temporal_Sup_L | Heschls_Gyrus_(includes_H1_and_H2) | 16.93 | 10.03 | 293.67 | 0.0020 | 0 |
| 122 | -63.6 | -22.8 | 8.9 | Temporal_Sup_L | Planum_Temporale | 8.52 | 7.93 | 340.62 | 0.0020 | 0 |
| 129 | -20.2 | -6.7 | -19.6 | Hippocampus_L | Left_Amygdala | 5.95 | 1.48 | 152.20 | 0.1080 | 0 |
| 130 | -22.7 | -6.4 | -20.4 | Amygdala_L | Left_Amygdala | 7.23 | 1.41 | 134.77 | 0.1290 | 0 |
| 131 | -25.2 | -5.9 | -21.3 | Amygdala_L | Left_Amygdala | 6.14 | 1.22 | 39.43 | 0.4110 | 0 |
| 132 | -27.6 | -5.3 | -22.4 | Amygdala_L | Left_Amygdala | 5.59 | 1.88 | 83.80 | 0.2610 | 0 |
| 133 | -30.1 | -4.9 | -23.6 | None | Left_Amygdala | 7.27 | 1.63 | 151.59 | 0.1020 | 0 |
| 137 | -27.5 | -21.5 | -19.9 | Hippocampus_L | Left_Hippocampus | 7.60 | 3.01 | 134.69 | 0.1440 | 0 |
| 138 | -32.9 | -22.2 | -20.5 | Fusiform_L | Left_Hippocampus | 5.87 | 6.73 | 232.31 | 0.0410 | 0 |
| 139 | -37.8 | -23.1 | -19.8 | Fusiform_L | Temporal_Fusiform_Cortex,_posterior_division | 9.24 | 5.17 | 586.96 | 0.0000 | 0 |
| 142 | -58.4 | -22.9 | -14.2 | Temporal_Mid_L | Middle_Temporal_Gyrus,_posterior_division | 10.52 | 4.90 | 468.97 | 0.0050 | 1 |
| 144 | -68.6 | -23.0 | -11.8 | Temporal_Mid_L | Middle_Temporal_Gyrus,_posterior_division | 8.42 | 1.84 | 161.00 | 0.0700 | 0 |
| 149 | -52.1 | -31.9 | -10.0 | Temporal_Mid_L | Middle_Temporal_Gyrus,_posterior_division | 5.05 | 9.03 | 518.56 | 0.0010 | 0 |
| 152 | -67.9 | -30.0 | -9.8 | Temporal_Mid_L | Middle_Temporal_Gyrus,_posterior_division | 9.77 | 4.47 | 333.65 | 0.0030 | 1 |

***:** If active TMS response > ± 5 SDs baseline, sham TMS response < ± 5 SDs baseline, and active TMS response was significantly stronger in nonparametric clustering analysis with 1000 permutations, then significance = 1 otherwise 0
